# Pharmacological activation of the circadian component REV-ERB inhibits HIV-1 replication

**DOI:** 10.1101/2020.04.09.031492

**Authors:** Helene Borrmann, Rhianna Davies, Matthew Dickinson, Isabela Pedroza-Pacheco, Mirjam Schilling, Alun Vaughan-Jackson, William James, Peter Balfe, Persephone Borrow, Jane A McKeating, Xiaodong Zhuang

## Abstract

Human immunodeficiency virus 1 (HIV-1) is a life-threatening pathogen that still lacks a curative therapy or vaccine. Despite the reduction in AIDS-related deaths achieved by current antiretroviral therapies, drawbacks including drug resistance and the failure to eradicate infection highlight the need to identify new pathways to target the infection. Circadian rhythms are endogenous 24-hour oscillations which regulate physiological processes including immune responses to infection, and there is an emerging role for the circadian components participating viral replication. The molecular clock consists of transcriptional/translational feedback loops that generate rhythms. In mammals, CLOCK and BMAL1 activate rhythmic transcription of genes including the nuclear receptor REV-ERBα, which represses BMAL1 and plays an essential role in sustaining a functional clock. We investigated whether REV-ERB activity regulates HIV-1 replication, and found REV-ERB agonists inhibited HIV-1 promoter activity in cell lines, primary human CD4 T cells and macrophages, whilst antagonism or genetic disruption of REV-ERB increased promoter activity. Furthermore, the REV-ERB agonist SR9009 inhibited promoter activity of different HIV-subtypes and HIV-1 replication in primary T cells. This study shows a role for REV-ERB synthetic ligands to inhibit HIV-1 LTR promoter activity and viral replication, supporting a role for circadian clock transcription factors in regulating HIV-1 replication.

## Introduction

All life forms have evolved under a rhythmically changing light/dark cycle due to the Earth’s rotation. From bacteria to man, all organisms possess an internal clock that oscillates in a 24-hour manner to anticipate environmental changes. The central clock and peripheral oscillators share a common molecular architecture and consist of transcriptional/translational feedback loops that regulate rhythmic gene expression [1]. In mammals, CLOCK and BMAL1 dimerize and the complex can bind E-box motifs in the promoter/enhancer of various clock genes, including *Per* and *Cry,* to activate their transcription. In turn, the PER and CRY proteins repress CLOCK/BMAL1 function and thereby shut down their own transcription. An additional feedback loop involves the nuclear receptors REV-ERBα and RORα. RORα competes with REV-ERBα for binding to the Bmal1 promoter ROR element (RORE) site and activates *Bmal1* transcription. REV-ERBα and RORα coordinate a regulatory loop which is crucial for stabilization of the core clock machinery [2] (Fig.1).

**Fig.1.**
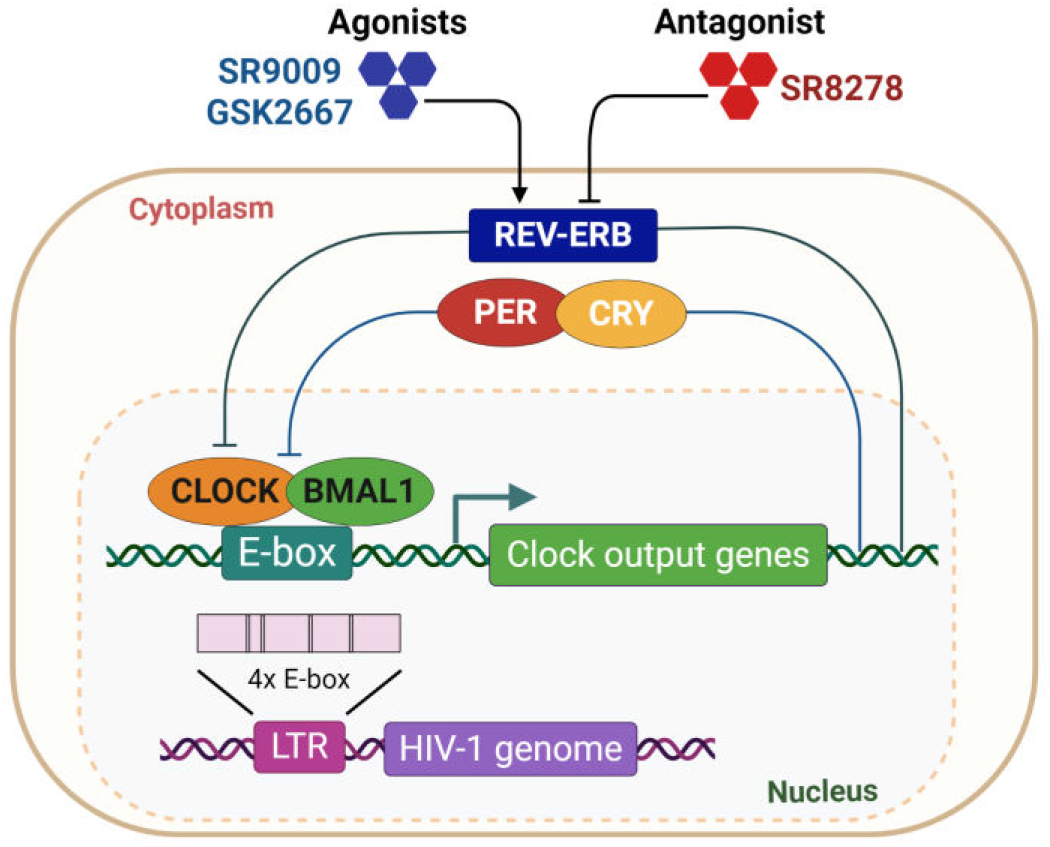
Schematic diagram illustrating the strategy for pharmacological modulation of REV-ERB.

The circadian system is recognized to play a role in regulating host innate and adaptive immune responses to microbial pathogens [3–5] and host susceptibility to an infectious agent is not only dependent on the inoculum size, transmission route and length of exposure, but on the time of day when the pathogen is encountered [6]. Recent clinical studies show that the time of vaccination can influence host immune responses and vaccine efficacy [7]. Viruses are obligate parasites that rely on host cell synthesis machinery for their replication, survival and dissemination. The potential for circadian pathways to regulate viral infection is an emerging research field [6, 8–10]. We recently reported a role for REV-ERB to regulate flavivirus replication and particle assembly, including hepatitis C virus, dengue virus and Zika virus [11].

Human immunodeficiency virus 1 (HIV-1) is the aetiologic agent of AIDS, one of the most devastating viral pandemics. Current therapies suppress HIV-1 replication and prevent the development of AIDS, but do not eradicate infection altogether. HIV-1 establishes latent sites of infection that promote viral persistence and evasion of host immune responses and anti-viral therapies [12]. HIV-1 primarily replicates in CD4 T cells and macrophages which display intrinsic rhythms of clock genes and cytokine expression [13–15]. Despite reports of disrupted circadian rhythms in HIV-1 infected patients [16–18], there is limited evidence supporting a direct role for circadian components in regulating HIV-1 replication. The HIV-1 long terminal repeat (LTR) promoter encodes regulatory elements that bind viral or cellular trans-activating factors that regulate its activity [19], demonstrating the innate dependency of the virus on host cell components to replicate. Chang et al. recently reported that BMAL1 positively regulates the HIV-1 LTR activity through E-box motifs therein [20] (Fig.1).

REV-ERBα/β are members of the nuclear hormone receptor family that are involved in the molecular clock circuits. Raghuram et al. identified haem as the physiological ligand of REV-ERB, showing that haem was required for recruiting the co-repressors: Nuclear Receptor CoRepressor (NCoR) and Histone Deacetylase 3 (HDAC3) [21]. Several synthetic ligands targeting REV-ERB have been developed, including the agonists (GSK4112 [22], SR9009 [23] and GSK2667 [24]) and antagonist (SR8278 [25]). Since REV-ERB can repress BMAL1 expression, these compounds are useful tools for examining circadian modulation and its effect on HIV-1 replication. In this paper, we show that REV-ERB synthetic ligands inhibit HIV-1 LTR promoter activity and viral replication, supporting a role for circadian clock transcription factors in regulating HIV-1 replication. This study highlights a novel research area with potential for discovery of new pathways that may impact on the replication of not only HIV-1, but also other viruses.

## Results and Discussion

A recent study reported that overexpression of BMAL1/CLOCK increased HIV-1 LTR activity [20], suggesting the potential for employing circadian modulators to inhibit BMAL1 and thereby reduce HIV-1 LTR activity. Since BMAL1 and CLOCK are basic helix-loop-helix PER-ARNT-SIM (bHLH-PAS) transcription factors, which are commonly considered undruggable, we tested the well characterized synthetic REV-ERB agonists SR9009 and GSK2667 and antagonist SR8278 for their ability to modulate HIV-1 LTR activity (Fig.1). Initial experiments were performed using the Hela TZM-bl cell line, which encodes integrated copies of the luciferase gene under the control of the HIV-1 LTR [26]. We confirmed these drugs had no cytotoxic effects on TZM-bl cells in the dose range and treatment length tested (SFig.1). Both REV-ERB agonists (SR9009 and GSK2667) inhibited basal HIV-1 LTR activity in a dose-dependent manner (Fig.2a–b), whilst the REV-ERB antagonist SR8278 increased LTR activity (Fig.2c). When treatment lengths of 4h, 8h and 24h were evaluated, the peak antiviral activity was seen after 24h of treatment for all three compounds (Fig.2d–f). To validate the specificity of the drugs, we treated TZM-bl cells with the agonists SR9009 or GSK2667 alone or in combination with antagonist SR8278. We observed a modest effect on basal LTR activity in the combined treatment group compared with sole treatment (Fig.2g–h), demonstrating that the compounds target a common molecular pathway.

**Fig.2.**
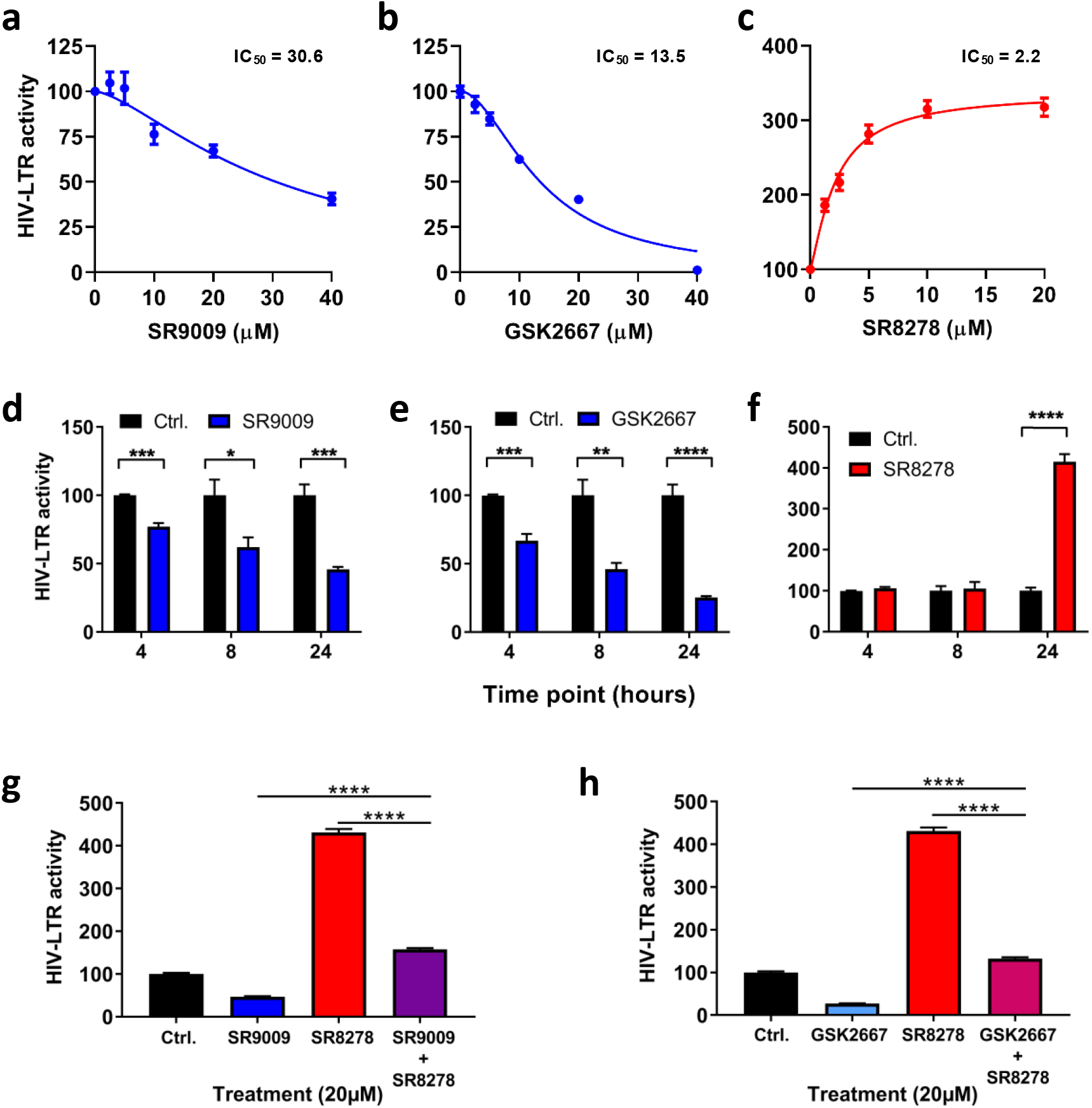
Pharmacological modulation of REV-ERB affects HIV-1 promoter activity. TZM-bl cells were treated with REV-ERB agonists SR9009 **(a)**, GSK2667 **(b)** or the antagonist SR8278 **(c)** at a range of doses for 24h and HIV-1 promoter activity measured by quantifying luciferase activity (mean ± S. E.M., n = 4). TZM-bl cells were treated with SR9009 **(d)** or GSK2667 **(e)** or SR8278 **(f)** at 20 μM for 4h, 8h or 24h and HIV-1 promoter activity measured by quantifying luciferase activity. Data are expressed relative to the control untreated cells at each time point (mean ± S.E.M., n = 4, Two-way ANOVA). **(g)** TZM-bl cells were treated with media containing SR9009 or SR8278, or simultaneously treated with SR9009 and SR8278 for 24 hours and HIV-1 promoter activity measured by quantifying luciferase activity (mean ± S.E.M., n = 8, One-way ANOVA). **(h)** TZM-bl cells were treated with GSK2667 or SR8278, or simultaneously treated with GSK2667 and SR8278 for 24 hours and HIV-1 promoter activity measured by quantifying luciferase activity (mean ± S.E.M., n = 8, One-way ANOVA). All dataare expressed relative to the control untreated cells.

Since the REV-ERB compounds target both REV-ERB isoforms, we investigated the effect of individual REV-ERBs on HIV-1-LTR activity. We silenced *Rev-erbα* or *Rev-erbβ* in TZM-bl cells using lentivectors expressing shRNA targeting either isoform. Since the lentivectors encoded a GFP reporter, the transduction efficiency was confirmed by imaging GFP expressing cells (Fig.3a). The knockdown efficiency was confirmed by qPCR of *Rev-erbα* or *Rev-erbβ* mRNA 48h post-transduction (Fig.3b). Silencing *Rev-erbα* or *Rev-erbβ* increased HIV-1 LTR activity (Fig.3c), suggesting a role for REV-ERBs in repressing HIV-1 promoter activity.

**Fig.3.**
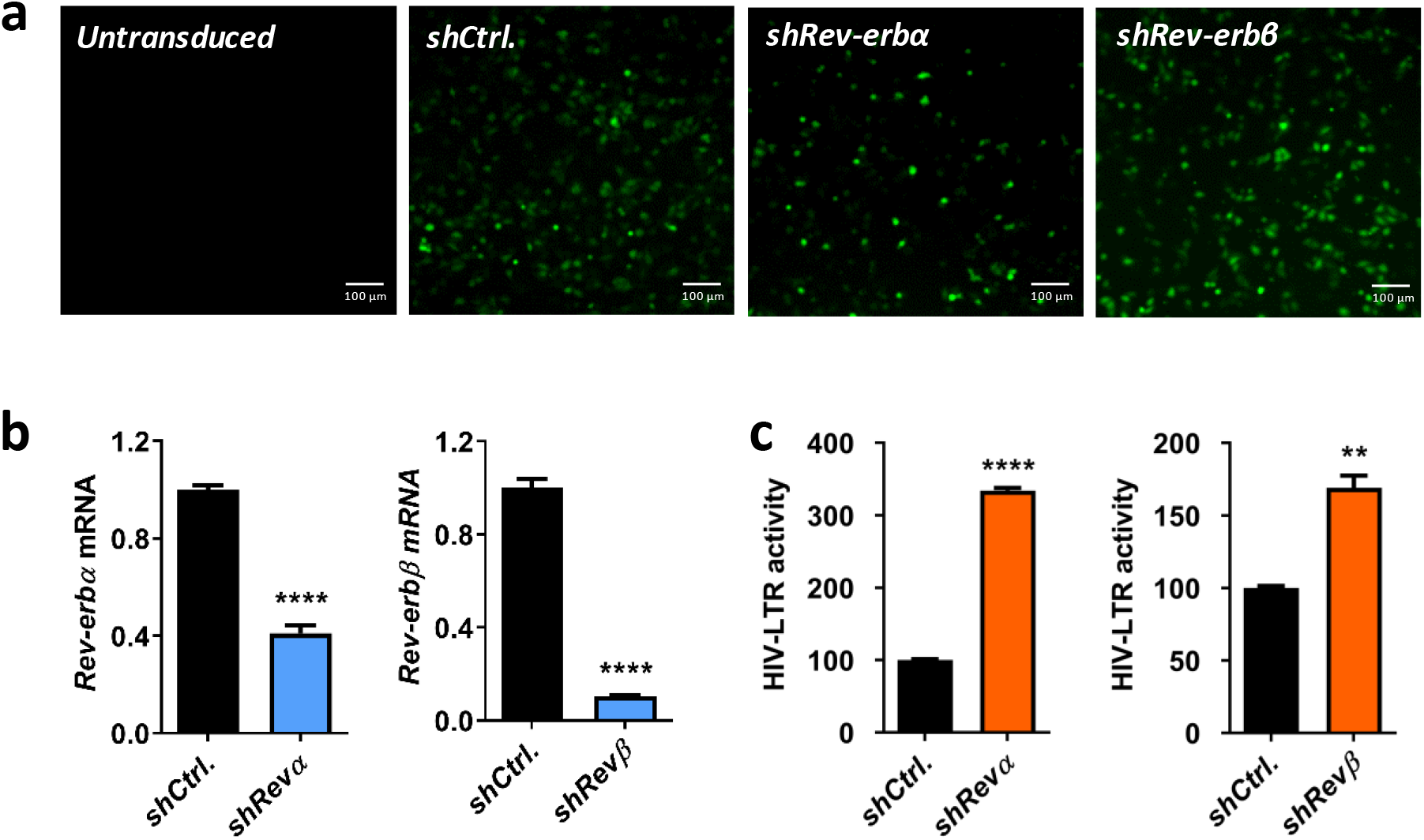
Silencing *Rev-erbs* increases HIV-1 promoter activity. **(a)** TZM-bl cells were transduced with control or *shRev-erbα* or *shRev-erbβ*-encoding lentiviral vectors that also expressed a GFP reporter, and fluorescent images of GFP were obtained 48h later to confirm transduction. **(b)** Total RNA was extracted from control or *Rev-erb* silenced TZM-bl cells at 48h post-transduction and *Rev-erbα* or *Rev-erbβ* mRNA levels quantified by qRT-PCR (mean ± S.E.M., n = 3, unpaired t test). **(c)** Control or *Rev-erb* silenced TZM-bl cells were lysed 48h post-transduction and HIV-1 promoter activity measured by quantifying luciferase activity (mean ± S.E.M., n = 3, unpaired t test).

To extend these observations to more physiologically relevant cell types, Jurkat cells (a CD4 T cell line) were infected with VSV-G complemented HIV NL4.3 R-E-reporter virus that undergoes a single cycle of replication that can be assessed by measuring luciferase activity. SR9009 significantly reduced HIV-1 replication and SR8278 showed an opposite effect (Fig.4a). Reassuringly, we show that these REV-ERB ligands regulate Bmal1 promoter activity in Jurkat cells (SFig.2). Importantly, these observations were replicated in activated primary human CD4 T cells and macrophages derived from human induced pluripotent stem cells (iPSC) (Fig.4b–c). Of note, these compounds showed no detectable cytotoxicity in any of the cell types used (SFig.3-5).

**Fig.4.**
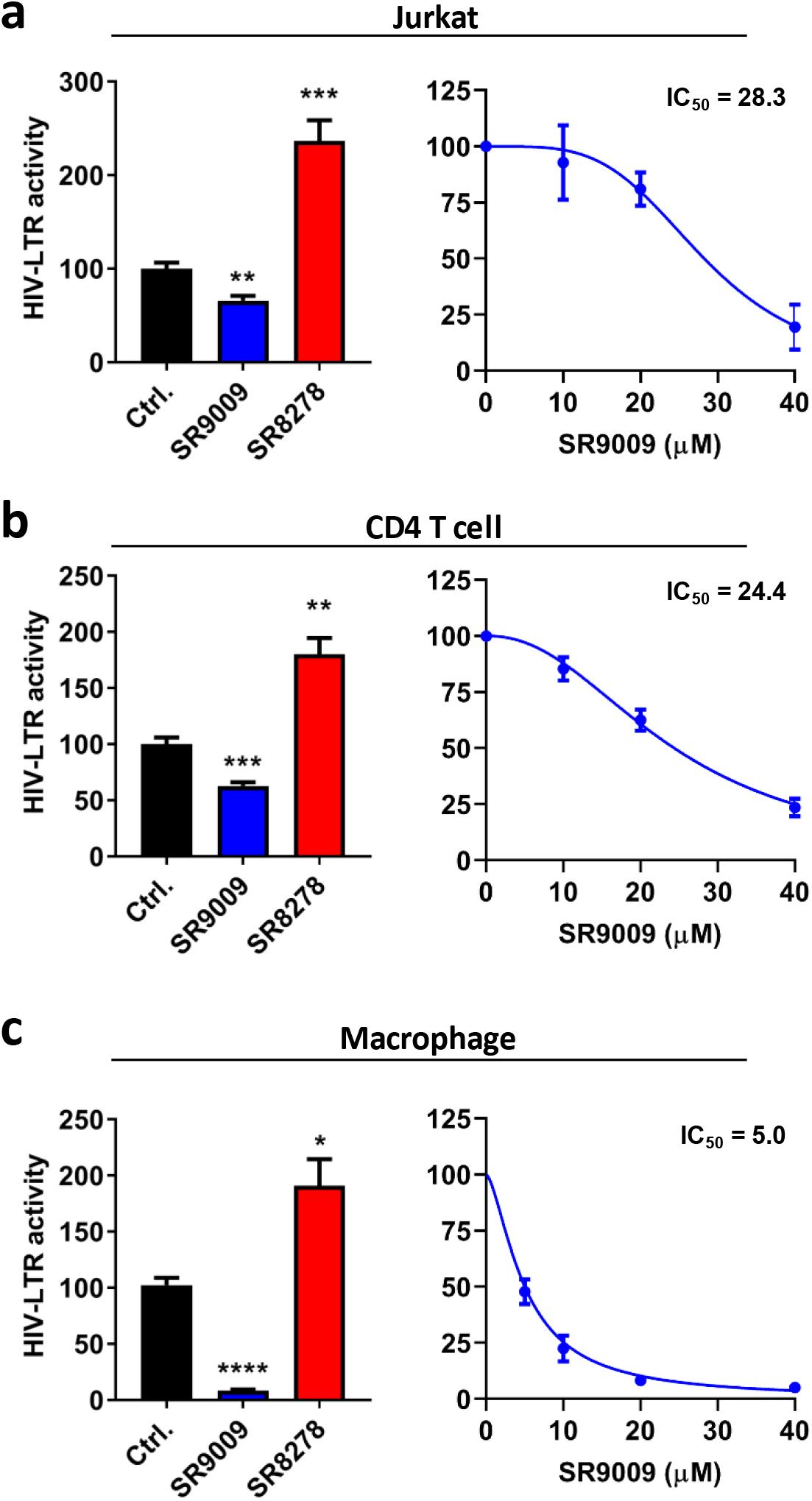
Pharmacological modulation of REV-ERB inhibits HIV-1 replication in T cells and macrophages. **(a)** Jurkat cells were infected with VSV-G-pseudotyped HIV-1 NL4.3-Luc for 24h. Infected cells were treated with SR9009 (20 μM) or SR8278 (20 μM) for 24h and LTR activity measured (mean ± S.E.M., n = 3, One-way ANOVA). The IC_50_ of SR9009 was determined at 24h post treatment by quantifying luciferase activity. **(b)** Primary CD4 T cells were activated for 3 days with anti-CD3/CD28 and infected with VSV-G-pseudotyped HIV-1 NL4.3-Luc for 24h and treated with SR9009 (20 μM) or SR8278 (10 μM) for 24h and LTR activity measured (n = 4, mean ± S.E.M, One-way ANOVA). The IC_50_ of SR9009 was determined at 24h post treatment by quantifying luciferase activity. **(c)** Human induced pluripotent stem cells (iPSCs) derived macrophages were infected with VSV-G-pseudotyped HIV-1 NL4.3-Luc for 24h and treated with SR9009 (10 μM) or SR8278 (5 μM) for 24h and LTR activity measured (n = 3, mean ± S.E.M, One-way ANOVA). The IC_50_ of SR9009 was determined at 24h post treatment by quantifying luciferase activity. All data are expressed relative to the control untreated cells.

The BMAL1/CLOCK complex binds a conserved motif (canonical CACGTG; non-canonical CANNTG) defined as the E-box and screening HIV-1 sequences deposited in the Los Alamos Database revealed that approximately one third of sequences (330/897) encode an E-box at position 294-286 in the U3 region of the LTR (Fig.5a). To investigate whether REV-ERB mediated-repression of the LTR was conserved across diverse HIV-1 subtypes, we transfected Jurkat cells with a panel of LTR-Luc reporter plasmids encoding promoter regions cloned from diverse HIV-1 clades [27]. All of the HIV-1 sub-type LTR-Luc reporters used in these experiments encode the canonical E-box motif (Fig.5b). As expected, SR9009 repressed the activity of all LTRs, demonstrating pan-subtype anti-viral activity (Fig.5b). Further analysis of these HIV-1 subtypes revealed additional conserved circadian regulatory elements in the LTR region, including the REV-ERB binding site RORE motif and the glucocorticoid response element (SFig.6). Cortisol production and secretion is tightly regulated by the circadian system and coordinates the synchronization of clock genes in peripheral tissues [28]. These data suggest a possibility where HIV-1 replication maybe synchronized in their reservoir cells at certain time of day. A recent study demonstrated a physical interaction of glucocorticoid receptor with REVERBα in the liver, providing an alternative mechanism for REV-ERB to regulate HIV-1 replication [29].

**Fig.5.**
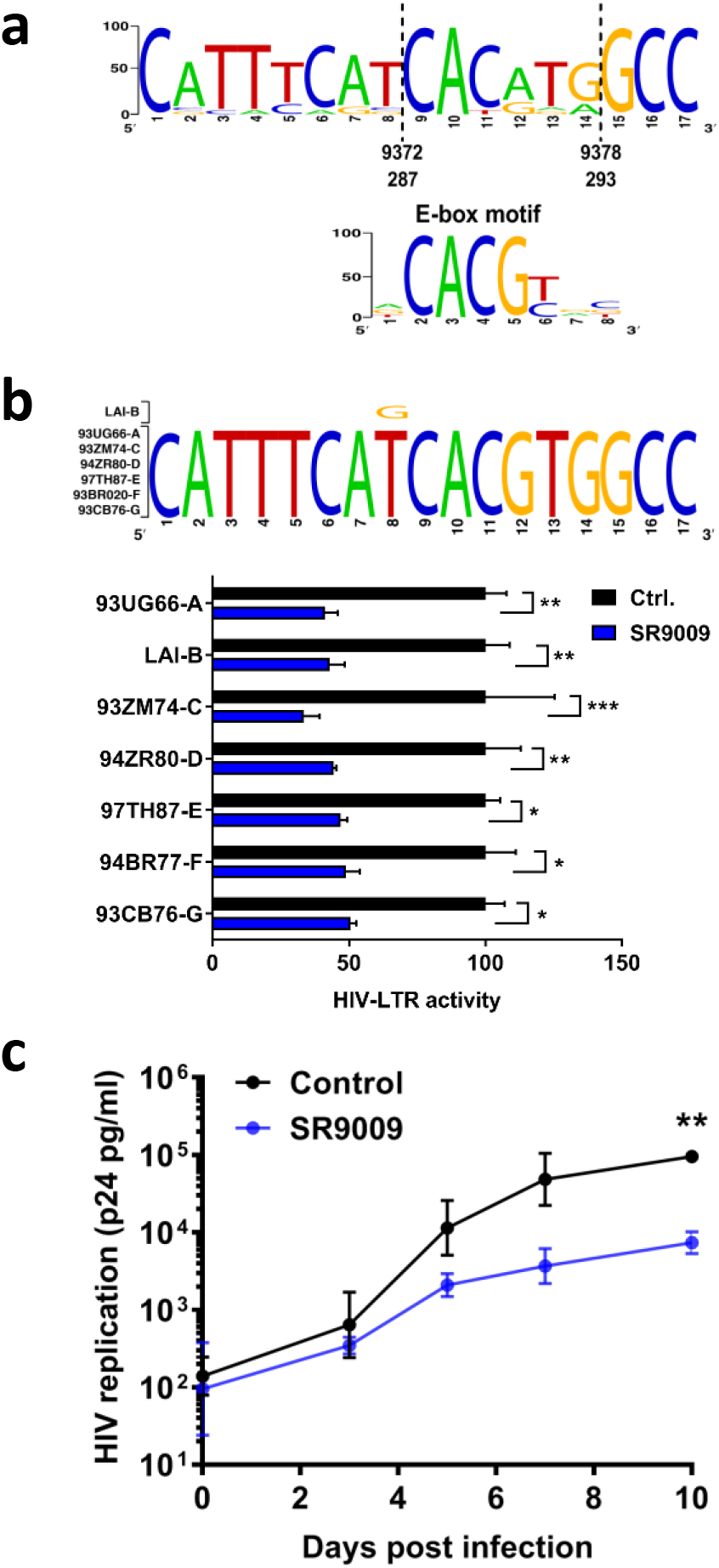
REV-ERB agonist SR9009 has pan-subtype antiviral activity. **(a)** Consensus plot illustrating the conserved nature of the E-box in the HIV-1 LTR based on the 897 HIV-1 sequences available in the LANL repository (analyzed with LANL QuikAlign/AnalyzeAlign software, coordinates are from the HXB2 referent). The level of conservation is reflected by the height of the bases (y axis 0 – 100%) [39, 40], with the consensus E-box motif (JASPAR database)[40] shown below. **(b)** Conserved E-box motifs in the HIV-1 strains chosen to study, where HIV LAI has a variant, G, at position 8. Jurkat cells were transfected with HIV-1 promoter reporter constructs for 24h. Transfected cells were treated with SR9009 at 20 μM for 24h and LTR activity measured by quantifying luciferase activity (mean ± S.E.M., n = 4, Two-way ANOVA). **(c)** CD8 depleted Human peripheral blood mononuclear cells (PBMCs) were infected with HIV-1 and cultured in medium with or without SR9009. Extracellular p24 levels were measured at intervals across a 10-day time period. Mean values from n = 3 biological repeats are shown, with error bars reflecting the geometric standard deviation. The first biological repeat employed cells from a single HIV-seronegative donor, whilst repeats two and three employed cells pooled from three healthy donors. The statistical significance of differences in mean extracellular p24 values between SR9009 and control cells at each time point was assessed using a paired t test.

Finally, we addressed the impact of SR9009 on the replication of an infectious virus in primary cells over time. Activated CD8 depleted peripheral blood mononuclear cells (PBMCs) from HIV-seronegative individuals were infected with a patient-derived subtype B virus (CH058) and cultured in medium with or without SR9009 (20 μM) for 10 days, reading out extracellular levels of p24 capsid antigen as a measure of viral replication at serial post-infection time points. While there was no impact on cell viability (SFig.7), SR9009 substantially reduced HIV-1 replication in this system, with a statistically significant difference in secreted p24 antigen levels observed by day 10 post infection (Fig.5c).

Our analysis of the E-box motifs in the HIV-1 LTR suggests the viral transcription is likely to be affected by the host circadian rhythm and more specifically by BMAL1. Cleret-Buhot et al. reported that CD4 Th17 cells are more permissive to HIV infection than other CD4 T subsets and showed increased *Bmal1* mRNA in CD4 Th17 lymphocytes [30], consistent with a positive regulatory role for BMAL1 in the HIV-1 replication cycle. One could speculate, since BMAL1 expression oscillates throughout 24h, HIV-1 may have evolved to adapt to this host rhythm by time of day dependent productive replication, to avoid host immune responses and to maximize viral persistence. Nevertheless our simplistic model system only demonstrated one aspect of the circadian regulation of HIV-1 which may under-represent the circadian impact if studied in a whole organism which warrants further investigations.

Since viruses are dependent on their host’s cellular machinery to replicate, it is likely that they have evolved to utilize host pathways to their advantage. In line with this, our data show that manipulating the host core circadian component REV-ERB can inhibit HIV-1 promoter activity. Importantly, our results suggest that REV-ERB ligands could be developed as a new class of HIV-1 replication inhibitors, which may synergize with current therapies. Of note, a recent report demonstrated REV-ERB dependent and independent effects of SR9009 [31], and therefore it is reassuring that *Rev-Erbα/β* silencing, along with an additional REV-ERB agonist GSK2667 and the antagonist SR8278 had consistent phenotypes to support a REV-ERB mediated effect. Our study highlights a novel and exciting area of HIV research and provides a rationale for future work to identify novel circadian modulators with anti-viral properties that may have potential to augment existing anti-viral regimens.

Current therapeutically-used combination anti-retroviral therapies effectively inhibit HIV-1 replication and delay progression of the disease. However viral resistance to antiviral agents can undermine their efficacy [32]. Instead of targeting the virus, the REV-ERB ligands target an intrinsic host pathway which is entirely different to the direct antiviral activity of the agents currently employed to block HIV-1 replication. REV-ERB agonists may provide an additional drug class as adjuvants or to aid in current combination therapy, especially in cases where there is a persistent low-level viraemia, or in patients where multi-class antiretroviral resistance is an obstacle to effective combination therapy. Moreover, the REV-ERB antagonist could have potential utility in ‘shock and kill’ HIV eradication strategies [33], as targeting this pathway may lead to fewer side effects than the agents currently being investigated in this approach.

## Methods

### Cell lines and primary cells

TZM-bl cells were provided by Professor Bill Paxton (University of Liverpool, Liverpool, UK) and cultured in DMEM medium supplemented with 10% heat-inactivated FBS (hiFBS,Sigma), 1% penicillin/streptomycin and 1% L-glutamine. Jurkat cells were provided by Professor Xiaoning Xu (Imperial College, London) and maintained in RPMI medium containing 10% hiFBS. Jurkat cells stably expressing Bmal1-luciferase promoter were generated using a well characterized lentiviral plasmid as previously described [34]. Peripheral blood mononuclear cells (PBMCs) were isolated from leukapheresis samples from healthy donors (NHS Blood and Transplant Service, Oxford, UK) and CD4 T cells isolated using the CD4+ T cell isolation kit from Miltenyi Biotec, Germany. Cells were stimulated with 50 IU/ml IL-2 (Proleukin; Novartis), 0.01ug/ml soluble human anti-CD3 (R&D clone UCHT1) and 0.1 ug/ml soluble human anti-CD28 (Life Technologies; clone CD28.2) at a concentration of 1×10^6^ cells/ml in RPMI-1640 (Life Technologies) containing 10% hiFBS, 1% penicillin/streptomycin (Sigma), 1% sodium pyruvate (Sigma), 1% Glutamax (Life Technologies), 1% non-essential aminoacids (Life Technologies) and 2 mM beta-mercaptoethanol (Life Technologies) (R10 medium), and incubated at 37°C in 5% CO_2_ for 3 days before infection. Human induced pluripotent stem cells (iPSC) derived macrophages were differentiated from the human iPSC line OX1-61 [35] and cultured in DMEM/F12 supplemented with 1% penicillin/streptomycin, Glutamax (2 mM), stabilized Insulin (5 μg/ml), HEPES pH 7.4 (15 mM), M-CSF (100 ng/ml) [36].

### Reagents

The following reagents were purchased from commercial suppliers: REV-ERB agonist SR9009 (Calbiochem, US); and REV-ERB antagonist SR8278 (Sigma, UK). REV-ERB agonist GSK2667 was synthesized at the University of Birmingham. All drugs were dissolved in dimethyl sulfoxide (DMSO) and their cytotoxicity determined using a lactate dehydrogenase (LDH) assay (Promega, UK).

### Flow cytometry

To measure cell viability, control or drug treated cells were stained with Live Dead Aqua (Life Technologies, UK). Cells were fixed with 4% PFA (Santa Cruz, UK) for 10 min at room temperature, then samples were acquired on a Cyan ADP flow cytometer (Beckman Coulter) and data analyzed using FlowJo (TreeStar).

### Plasmids

Lenti-shRev-Erbα/β constructs were gifts from Dr. B. Grimaldi, University of Genoa, Italy [37]. The plasmid encoding HIV NL4.3 R-E-Luciferase was obtained from the NIBSC AIDS Repository. VSV-G expression plasmid was previously reported [38]. The single-cycle HIV-1 pseudotyped with the VSV-G envelope were generated in 293T cells using the HIV NL4.3 R-E-Luciferase plasmid. Plasmids encoding full-length infectious molecular clones of NL4.3-Bal were obtained from Drs John Kappes and Christina Ochsenbauer-Jambor (University of Alabama at Birmingham, US). HIV-LTR Luc constructs were previously reported [27] and provided by Professor Bill Paxton (University of Liverpool, Liverpool, UK).

### HIV LTR E-box analysis and subtypes promoter activity

The HIV-1 LTR sequences deposited in the Los Alamos Database were searched for the E-box motif CACGTG using the Sequence Search Interface program (www.hiv.lanl.gov) and the number and location of matches enumerated. Consensus plot illustrating the conserved nature of the E-box in the HIV-1 LTR based on the 897 HIV-1 sequences available in the LANL repository [39, 40]. HIV-1 LTR-Luc plasmids were delivered into Jurkat cells using the Neon^®^ Transfection System following the manufacturer's instructions. Luciferase activity was measured using the Promega kit following the manufacturers’ instructions (Promega, UK).

### Generation and titration of viral stocks

To produce HIV CH058 6M stocks, plasmid DNA was transfected into 293FT cells using Lipofectamine (Life Technologies, UK) or Fugene 6 (Promega, UK) and virus containing supernatants harvested 3 days later, clarified by centrifugation at 1400 x g for 10 min and stored at −80°C. The infectivity of viral stocks was determined using a colorimetric reverse transcriptase assay (Roche Life Sciences). VSV-G complemented NL4.3 R-E-viral stocks were generated as previously reported [38] and RT activity measured using a PCR based method [41].

### In vitro HIV-1 replication assay

PBMCs from HIV-seronegative donors were depleted of CD8+ T cells using CD8 microbeads (Miltenyi Biotec, Germany). Cells from a single donor or pooled cells from 3 different donors were activated by culturing for 3 days in R10 medium with 50IU/ml IL-2 and antibodies to CD3 and CD28 as detailed above. Cells were then plated at 200,000 cells/well into 96-well round-bottomed microplates in triplicate, and infected by spinoculation with HIV-1 derived from an infectious molecular clone corresponding to the consensus quasispecies sequence of the virus present in HIV-1 infected patient CH058 at 6 months post-infection (CH058 6M) [42] at a concentration of 0.25ng reverse transcriptase/10^6^ cells. After a two-hour spinoculation, cells were washed twice before culturing in R10 medium with 50IU/ml IL-2 with or without SR9009 (20 μM). Supernatants were harvested 90 minutes later (“day 0”) and at two to three day intervals thereafter for a 10-day period and stored at −80°C. Supernatants were subsequently analysed for HIV-1 capsid antigen using a p24 alphaLISA^®^ assay (Perkin-Elmer, US). The assay was performed according to the manufacturer’s instructions, and plates were read on FLUOstar Omega plate reader (BMG Labtech, Germany).

### Statistical Analysis

All experiments were performed at least twice. All data are presented as mean values ± SEM. P values were determined using the unpaired t test (two group comparisons; unpaired data) or paired t test (two group comparison; paired data) using PRISM version 8. In the figures ∗ denotes p < 0.05, ∗∗denotes p < 0.01, ***denotes p <0.001, ****denotes p <0.0001, n.s. denotes non-significant.

## Supporting information

SUPPLEMENTARY MATERIALS for MS: Pharmacological activation of the circadian component REV-ERB inhibits HIV-1 replication

## Data availability statement

All data generated or analysed during this study are included in this published article (and its Supplementary Information files).

## Acknowledgements

We thank Bill Paxton (University of Liverpool) for diverse HIV-LTR Luc plasmids and TZM-bl cells and Xiaoning Xu (Imperial College, London) for Jurkat cells. This work was funded by Wellcome Trust award IA 200838/Z/16/Z (JAM); MRC project grant MR/R022011/1 (JAM) and NIH, NIAID, DAIDS grants R01 AI 114266 and UM1 AI 126619 (P Borrow). P Borrow is a Jenner Institute Investigator.

## Author contributions

HB conducted experiments; RD conducted experiments; MD conducted experiments; IP-P, MS, AVJ and WJ provided reagents; P Balfe performed data analysis; P Borrow contributed to experimental design, provided critical comments and edited the MS; JAM designed study and co-wrote MS; XZ designed study, conducted experiments and wrote the MS.

