## SUPPLEMENTARY MATERIALS for MS: Pharmacological activation of the circadian component REV-ERB inhibits HIV-1 replication for "Pharmacological activation of the circadian component REV-ERB inhibits HIV-1 replication"

*Shared first authorship

### Supplementary figure 1

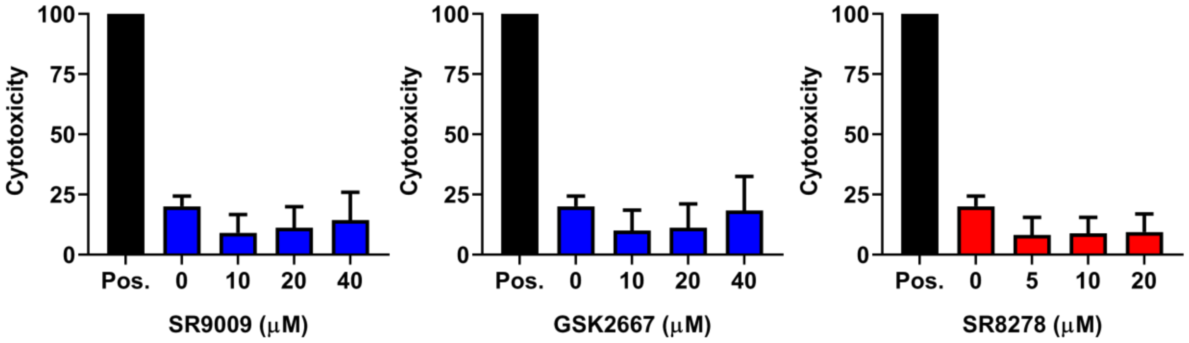


**SFig.1 Lack of cytotoxicity of REV-ERB ligands for TZM-bl cells.** TZM-bl cells were treated with REV-ERB agonists SR9009 or GSK2667 or the antagonist SR8278 at a range of doses for 24h and cytotoxicity determined using an LDH assay (mean ± S.E.M., n = 2). Data are expressed relative to the positive control representing total cell lysate (100% cytotoxicity).

#### Supplementary figure 2

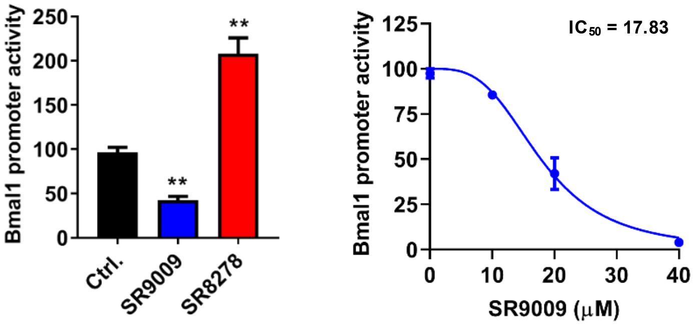


**SFig.2 Effect of REV-ERB ligands on Bmal1 promoter activity in Jurkat cells.** Jurkat cells stably expressing a Bmal1 promoter-luciferase construct were treated with the REV-ERB agonist SR9009 (20 μM) or antagonist SR8278 (20 μM) and luciferase activity measured after 24h (mean ± S.E.M., n = 3, One-way ANOVA). The $IC_{50}$ of SR9009 was determined at 24h post treatment by quantifying luciferase activity.

### Supplementary figure 3

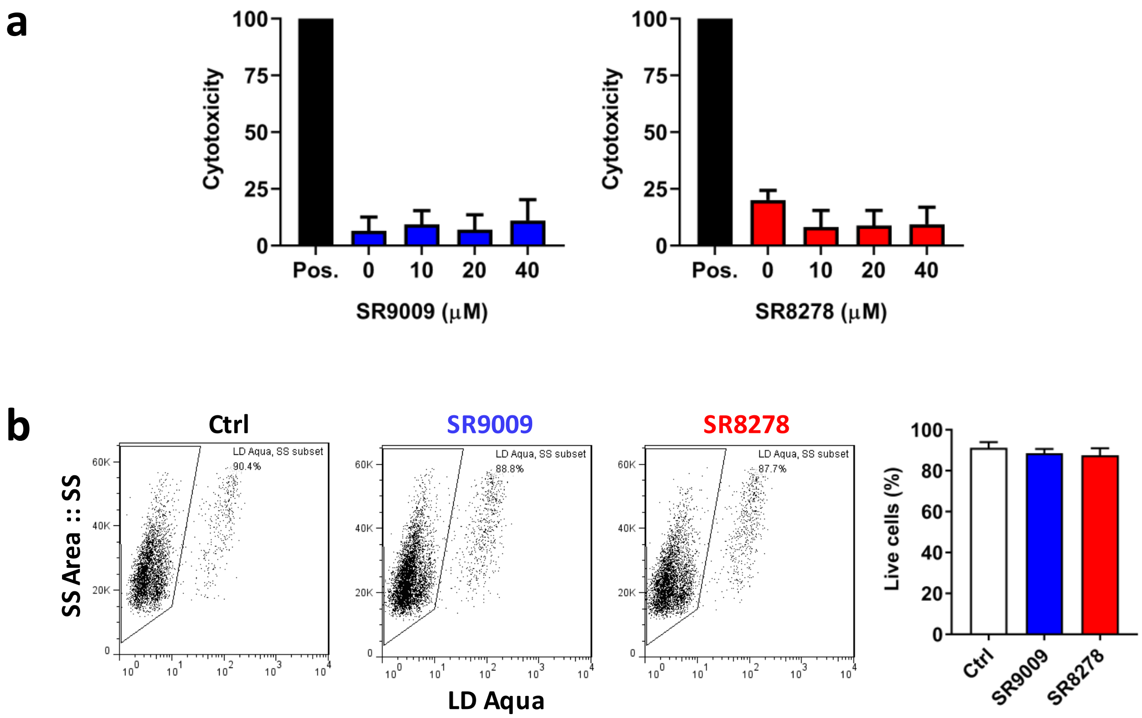


**SFig.3 Lack of cytotoxicity of REV-ERB ligands for Jurkat cells. (a)** Jurkat cells were treated with the REV-ERB agonist SR9009 or antagonist SR8278 at a range of doses for 24h and cytotoxicity determined using an LDH assay (mean ± S.E.M., n = 2). Data are expressed relative to a positive control representing total cell lysate (100% cytotoxicity). **(b)** Jurkat cells were treated with the REV-ERB agonist SR9009 or antagonist SR8278 for 24h or medium only as a negative control, and viability assessed by flow cytometry using a live-dead stain. Dot plots illustrate staining of one representative biological sample and summary plots of the percentage of live cells are shown on the right (n=7, mean + S.E.M., One-way ANOVA).

### Supplementary figure 4

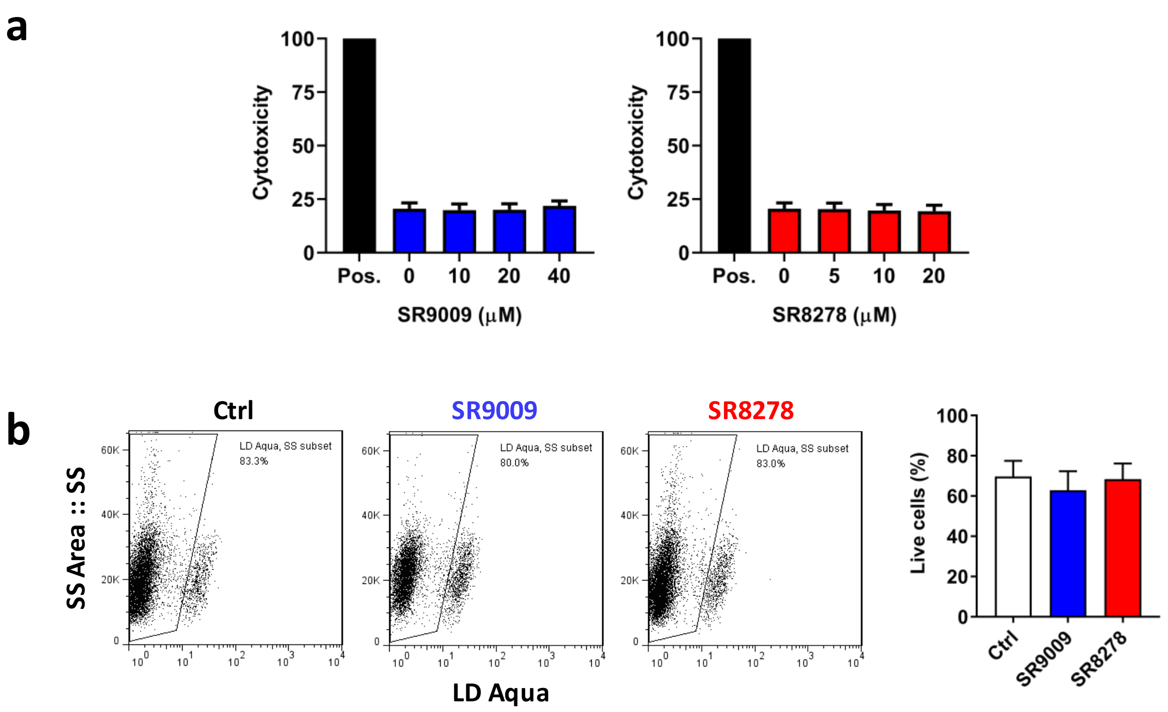


**SFig.4 Lack of cytotoxicity of REV-ERB ligands for human primary CD4 T cells.** Primary human CD4 T cells were activated with anti-CD3/CD28 for 3 days and treated with the REV-ERB agonist SR9009 or antagonist SR8278 at a range of doses for 24h and cytotoxicity determined using an LDH assay (mean $\pm$ S.E.M., n = 3). Data are expressed relative to the positive control representing total cell lysate (100% cytotoxicity). **(b)** Activated CD4 T cells were treated with the REV-ERB agonist SR9009 or antagonist SR8278 for 24h or medium only as a negative control, and cell viability assessed by flow cytometry using a live-dead stain. Dot plots illustrates staining of one representative biological sample and summary plots of the percentage of live cells are shown on the right (n=7, mean + S.E.M., One-way ANOVA).

### Supplementary figure 5

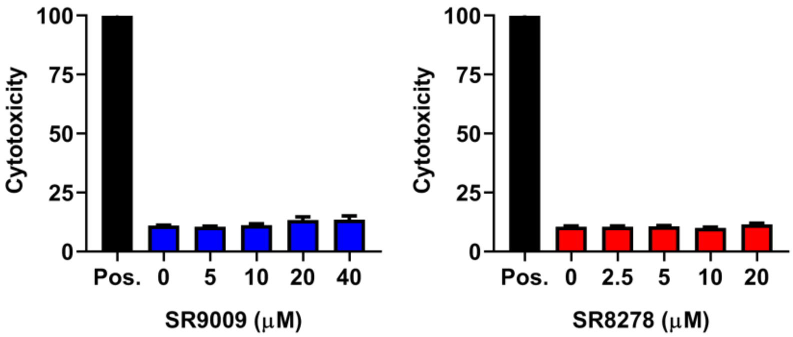


**SFig.5 Lack of cytotoxicity of REV-ERB ligands for human induced pluripotent stem cells (iPSCs) derived macrophages.** Human iPSCs-derived macrophages were treated with the REV-ERB agonist SR9009 or antagonist SR8278 at a range of doses for 24h and cytotoxicity determined using a LDH assay (mean ± S.E.M., n = 3). Data are expressed relative to the positive control representing total cell lysate (100% cytotoxicity).

### Supplementary figure 6

**ROR response element (Start 412 bp)**

```
          **********************
LAI-B     CCATCCAAAGGTCAGTGGATAT
93UG66-A  CCATCCAAAGGTCAGTGGATAT
93ZM74-C  CCATCCAAAGGTCAGTGGATAT
94ZR80-D  CCATCCAAAGGTCAGTGGATAT
97TH87-E  CCATCCAAAGGTCAGTGGATAT
93BR020-F CCATCCAAAGGTCAGTGGATAT
93CB76-G  CCATCCAAAGGTCAGTGGATAT
```

**Glucocorticoid receptor elements (start: 343 bp)**

```
          **********************
LAI-B     CAAGCTGGTGTTCTCTCCTTTA
93UG66-A  AAAGCTGGTGTTCTCTCCTTTA
93ZM74-C  CAAGCTGGTGTTCTCTCCTTTA
94ZR80-D  CAAGCTGGTGTTCTCTCCTTTA
97TH87-E  CAAGCTGGTGTTCTCTCCTTTA
93BR020-F CAAGCTGGTGTTCTCTCCTTTA
93CB76-G  CAAGCTGGTGTTCTCTCCTTTA
```

**SFig.6 Conserved ROR response element and glucocorticoid receptor element in the HIV-1 LTR.**

### Supplementary figure 7

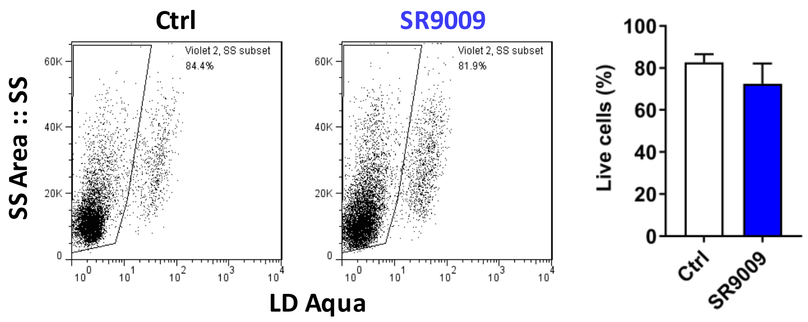


**SFig.7 Viability of peripheral blood mononuclear cells (PBMCs) following extended treatment with SR9009.** Human PBMCs were depleted of CD8 T cells and activated by culture with antibodies to CD3 and CD28 for 3 days, then infected with HIV-1 and cultured in medium with or without SR9009 (20 µM) for a further 7 days. The first biological repeat employed cells from a single HIV-seronegative donor, whilst repeats two and three employed cells pooled from three healthy donors. Cell viability was assessed by flow cytometry using a live-dead stain on day 7 post SR9009 treatment. Dot plots illustrate staining of one representative biological sample and summary plots of the percentage of live cells are shown on the right (n=3, mean + S.E.M., paired t test).
